# Selective advantages favour high genomic AT-contents in intracellular elements

**DOI:** 10.1101/448324

**Authors:** Anne-Kathrin Dietel, Holger Merker, Martin Kaltenpoth, Christian Kost

## Abstract

Extrachromosomal genetic elements generally exhibit increased AT-contents relative to their hosts’ DNA. The AT-bias of endosymbiotic genomes is commonly explained by neutral evolutionary processes. Here we show experimentally that an increased AT-content of host-dependent elements can be selectively favoured on the host level. Manipulating the nucleotide composition of bacterial cells by introducing A+T-or G+C-rich plasmids, we demonstrate that cells containing GC-rich plasmids are less fit than cells containing AT-rich plasmids. Moreover, the cost of GC-rich elements could be compensated by providing G+C-, but not A+T-precursors, thus linking the observed fitness effects to the cytoplasmic availability of nucleotides. Our work identifies selection as a strong evolutionary force that drives the genomes of intracellular genetic elements toward higher A+T contents.

**Author Summary:** Genomes of endosymbiotic bacteria are commonly more AT-rich than the ones of their free-living relatives. Interestingly, genomes of other intracellular elements like plasmids or bacteriophages also tend to be richer in AT than the genomes of their hosts. The AT-bias of endosymbiotic genomes is commonly explained by neutral evolutionary processes. However, since A+T nucleotides are both more abundant and energetically less expensive than G+C nucleotides, an alternative explanation is that selective advantages drive the nucleotide composition of intracellular elements. Here we provide strong experimental evidence that intracellular elements, whose genome is more AT-rich than the genome of the host, are selectively favored on the host level. Thus, our results emphasize the importance of selection for shaping the DNA base composition of extrachromosomal genetic elements.

## Introduction

Bacterial genomes exhibit a considerable amount of variation in their nucleotide composition (G+C versus A+T), ranging from less than 13% to more than 75% GC (1, 2). Despite intense efforts during the past decades, the selective pressures determining the evolution and maintenance of this variation remain elusive. A general pattern that emerged from sequencing the genomes of numerous taxa is that bacteria, whose survival obligately depends on a eukaryotic host (i.e. endosymbionts), display genomic AT-contents that are significantly increased in comparison to the genomes of their free-living relatives as well as their hosts’ genomes (3). Interestingly, intracellular genetic elements that permanently or transiently exist outside the bacterial chromosome, such as plasmids, viruses, phages, and insertion sequence (IS) elements, are also usually characterized by a significantly higher AT-content than the genome of their host (4, 5). Finding that the nucleotide composition of these very different elements is consistently biased in the same direction suggests similar evolutionary mechanisms operate to produce this pattern.

While less attention has been paid to extrachromosomal genetic elements such as plasmids and bacteriophages, two main hypotheses have been put forward to explain the biased nucleotide composition of obligate intracellular bacteria: First, high AT-contents can result from increased levels of genetic drift and mutational bias (3, 6, 7). Genetic drift is particularly strong when the bacteria’s effective population sizes are small, which is generally the case in vertically transmitted, intracellular symbionts (3). Since the majority of DNA modifications caused by oxygen radicals (either from the environment or generated by endogenous cellular processes) lead to mispairing of DNA bases, which mostly results in GC→AT transitions and G/C→T/A transversions (8), in the long-run genetic drift is expected to increase the elements’ overall AT-content.

On the other hand, it has been argued that the AT-bias of intracellular elements could be adaptive, and thus favored by natural selection (4). The reasoning behind this idea is that both endosymbiotic bacteria and plasmids occupy the same ecological niche, i.e. the intracellular environment of a larger organism, and thus have access to metabolites in the host’s cytoplasm (9). This includes all nucleotides and their biochemical precursors. For the host cell, ATP and UTP nucleotides are energetically less expensive to produce than GTP and CTP nucleotides (4). Moreover, ATP is the main energy currency used in cells and, thus, the most abundant nucleotide (10, 11). Hence, a preferential uptake of A+T nucleotides by the respective intracellular element may impose a lower metabolic burden on its host than consumption of the more valuable G+C nucleotides. Strikingly, in both endosymbiotic bacteria and plasmids, selection tends to reduce the costs intracellular elements impose on their host (e.g. by reducing the size and transcriptional activity of the element (9)). The reason for this is that hosts harbouring metabolically ‘costly’ intracellular elements display a lower fitness than hosts with metabolically ‘cheaper’ elements (12). As a consequence, selection acts against the less fit symbiont-host combinations, thereby favouring hosts that harbour metabolically cheaper intracellular elements (host-level selection). Accordingly, if AT-rich elements are metabolically ‘cheaper’ than GC-rich elements, hosts with more AT-rich elements should be selectively favoured.

Until now, GC-content variation in bacteria has mainly been studied using comparative approaches (13, 14) (but see (15)). This is likely due to technical difficulties to experimentally disentangle and manipulate these complex and often obligate host-endosymbiont systems (16). Although understandable from a methodological point of view, sequence comparisons can only reveal correlative relationships. To identify the underlying mechanistic causes, however, manipulative experiments are required that allow to rigorously scrutinize the focal hypothesis under controlled conditions.

While the GC-content evolution in plasmids has also mainly be studied using comparative approaches, plasmids and their hosts are highly tractable systems, in which a variety of different features can be experimentally manipulated (9). Here we take advantage of the experimental tractability of plasmid-host interactions to unravel the evolutionary consequences resulting from a biased nucleotide composition of host-dependent, intracellular elements. Manipulating the plasmids’ GC-content allowed us to experimentally test the hypothesis that a higher demand for GC-nucleotides due to the presence of a symbiont (here: plasmid) limits host growth. Our results show indeed that bacteria containing GC-rich plasmids and thus having an increased demand for G+C-nucleotides were less fit than bacteria with AT-rich plasmids. Supplying cells that contained a GC-rich plasmid with G+C nucleosides restored the fitness of host cells, while no such effect was observed for cells with AT-rich plasmids or supplementation with A+T nucleotides to both types of plasmid-containing cells. These findings suggest that the cytoplasmic availability of G+C nucleotides limited the growth of cells with GC-rich plasmids. Moreover, introducing plasmids into different bacterial species with an increasing genomic GC-content revealed that fitness costs imposed by GC-rich plasmids decreased with increasing GC content of the host genome. Taken together, our results provide strong experimental evidence that the commonly observed increased AT-content of host-dependent elements can be selectively favoured.

## Results

### Experimental model system

To determine if differences in the nucleotide composition of an extrachromosomal genetic element affect the fitness of the corresponding host cell, plasmids with high or low GC-contents were introduced into *Escherichia coli* cells. For this, two plasmids served as a backbone, into which eight non-coding AT-or GC-rich sequences of 1 kb in size were introduced to alter their net GC-content (Fig. 1). Sequences originated from eukaryotic DNA and were carefully selected, such that no genes or regulatory elements were present (see Materials and Methods). In this way, chances of inadvertent gene expression were minimized, which could have resulted in additional metabolic costs. All AT-and GC-rich sequences were individually introduced into two different plasmid backbones that strongly differ in terms of their genetic architecture (i.e. origin of replication, copy number, and selectable marker). The resulting plasmid constructs (i.e. eight AT-rich and eight GC-rich plasmids for each of the two backbones) were used as replicates to rule out plasmid-or sequence-specific effects. This system allowed us to study the fitness consequences resulting from intracellular elements that differed in their genomic nucleotide composition in otherwise isogenic bacterial cells.

**Fig.1.**
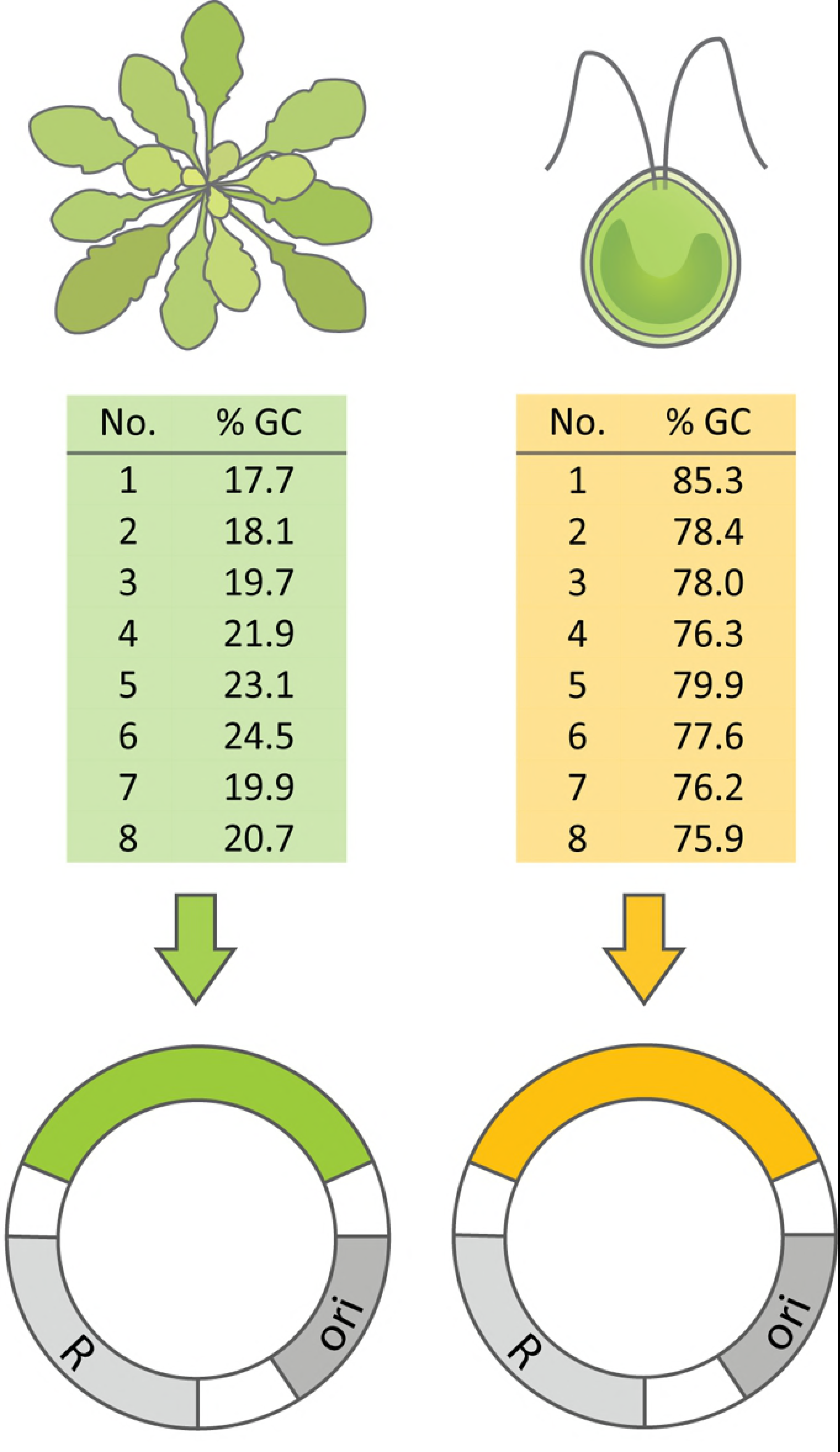
Construction of AT-and GC-rich plasmids. Eight non-coding AT-and GC-rich sequences of 1 kb in size were amplified from the genomes of *Arabidopsis thaliana* and *Chlamydomonas reinhardtii,* respectively. Sequences were inserted into two different plasmid backbones (pJet and pBAV) to exclude plasmid-specific effects. Different plasmid-insert combinations were treated as independent replicates. All plasmids contained an origin of replication (ori), a resistance cassette (R), and one of the inserts (AT-rich DNA: green label, GC-rich DNA: orange label).

### *E. coli* cells containing AT-rich plasmids are fitter than cells containing GC-rich plasmids

To determine whether the plasmids’ nucleotide composition affected the host cell’s fitness, AT-rich (i.e. cells harbouring AT-rich plasmids) and GC-rich (i.e. cells harbouring GC-rich plasmids) *E. coli* cells were grown for 24 h in minimal medium, and the growth kinetics of these cultures were determined spectrophotometrically. The results of this experiment revealed that GC-rich cells harbouring the plasmid pJet1.2/blunt (hereafter: pJet) grew significantly less well than the corresponding AT-rich cells: GC-rich cells displayed a significantly extended lag-phase, a decreased maximum growth rate, and reached a lower maximum cell density as compared to AT-rich cells (Fig. 2 A-C). Repeating the same experiments with *E. coli* strains that harboured the second plasmid backbone (i.e. pBAV1kT5gfp (17) lacking *T5gfp;* hereafter: pBAV), corroborated these results. Again, cell populations containing GC-rich plasmids displayed a significantly extended lag-phase, a decreased maximum growth rate, and reached a significantly reduced maximum cell density relative to the cognate AT-rich cells (Fig. 2 D-F). Taken together, GC-rich cells grew significantly less well than AT-rich cells, indicating fitness costs resulted from the presence of GC-rich plasmids. Due to the design of the experiment, effects that might have been caused by the inserted sequences or the plasmid backbones used could be ruled out as explanation.

**Fig.2.**
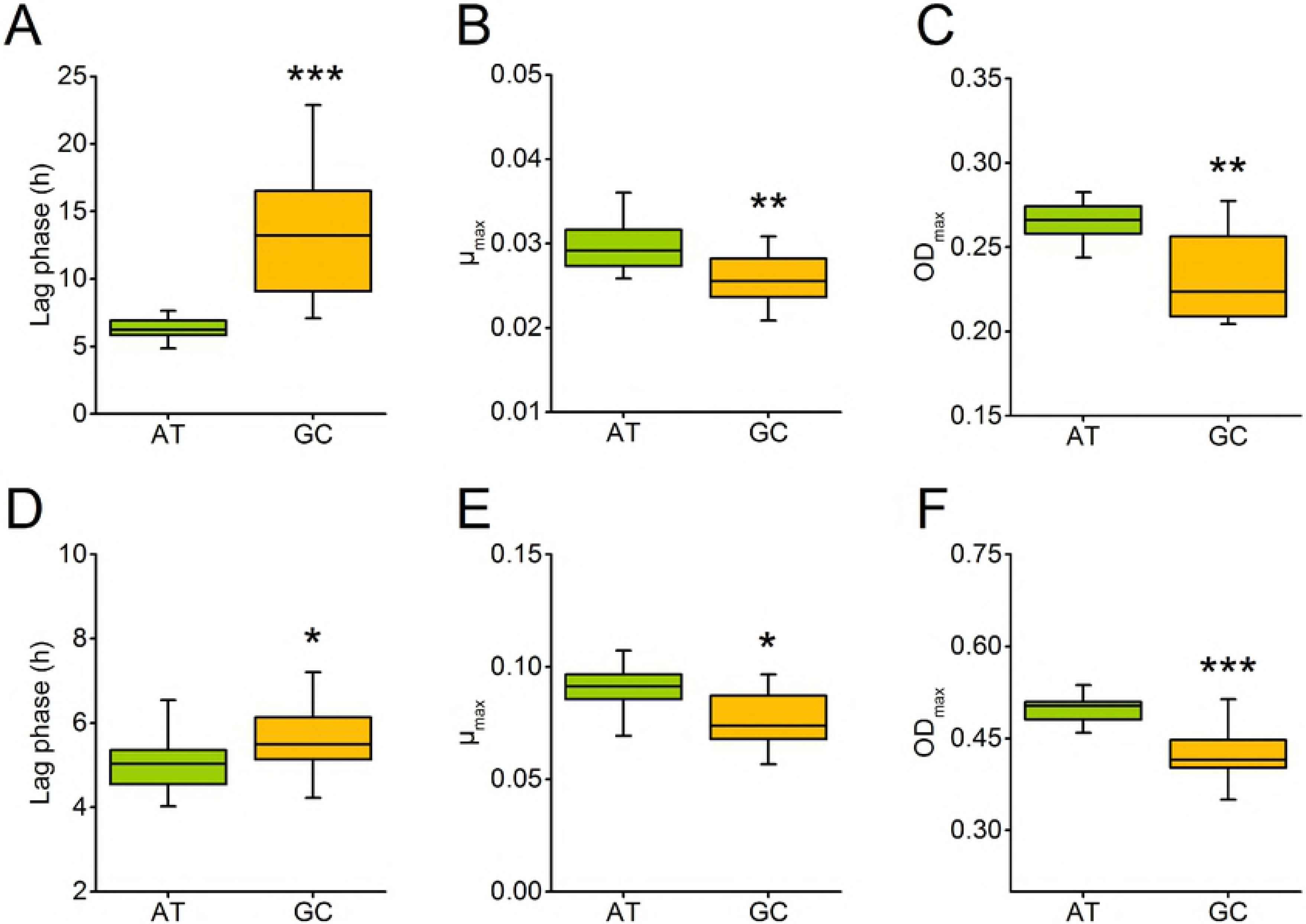
Cells containing GC-rich plasmids are less fit than those with AT-rich plasmids. Growth experiments of *E. coli* harbouring AT-(green) or GC-rich (orange) pJet (**A,B,C**) or pBAV plasmids (**D,E,F**) were performed in minimal medium. Growth over 24 h measured as optical density at 600 nm was used to calculate fitness-relevant parameters. (**A,D**) Duration of lag phase, (**B,E**) maximum growth rate, and (**C,F**) maximum optical density reached after 24 h of cells harbouring either AT-rich or GC-rich plasmids. Asterisks indicate significant differences between cells containing AT-and GC-rich plasmids. Independent-samples t-test: *** P < 0.001, ** P < 0.01, and * P < 0. 05, n = 16.

### GC-rich plasmids have a lower copy number than AT-rich plasmids

The copy number of plasmids is usually genetically determined by replication-control mechanisms that are encoded by the plasmid itself (18). Nevertheless, the copy number of a single plasmid can vary, depending on the extent of metabolic costs it imposes on its host. For example, the copy number of a plasmid has been shown to decrease with increasing length or gene content of the accessory region (12). Accordingly, if a GC-rich plasmid imposes a higher metabolic cost on its host than an AT-rich plasmid, it should also be present in a lower copy number than the less costly AT-rich plasmid. This hypothesis was tested by quantifying the copy numbers of both plasmids harbouring AT-or GC-inserts via quantitative PCR (qPCR). Indeed, qPCR analyses revealed that the copy number of both plasmid backbones analysed were drastically reduced when plasmids contained GC-rich inserts relative to plasmids with AT-rich inserts (Fig. 3). These results further corroborate that GC-rich plasmids likely impose a higher metabolic burden on their host cells than AT-rich plasmids, thus resulting in a strongly reduced copy number.

**Fig.3.**
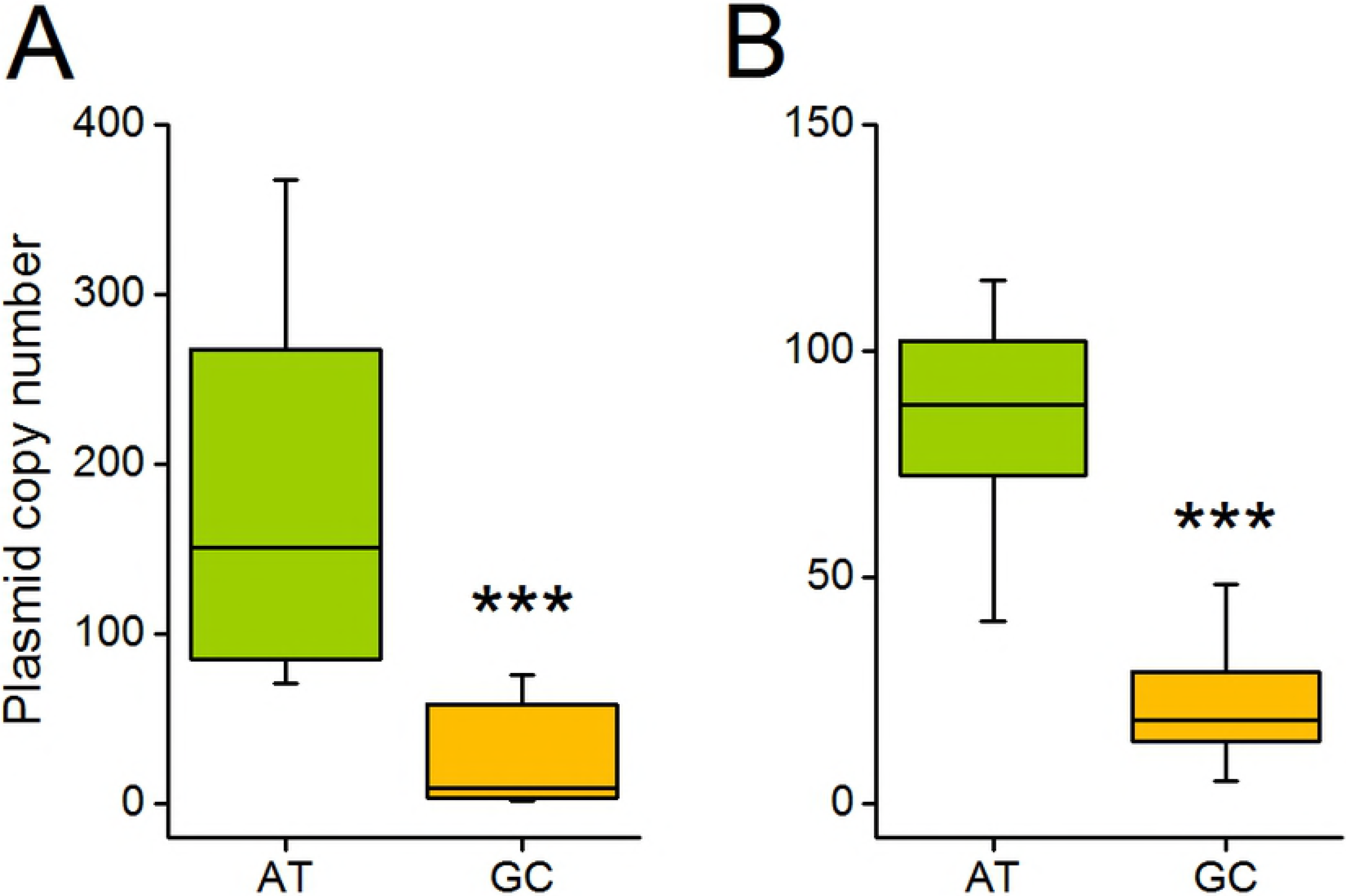
GC-rich plasmids have a lower copy number than AT-rich plasmids. Plasmid copy number per bacterial chromosome equivalent was assessed by quantitative realtime PCR of *E. coli* cultures after 24 h of growth. Shown are copy numbers of AT-(green) and GC-rich (orange) (**A**) pJet plasmids and (**B**) pBAV plasmids. Independent-samples t-test: *** P < 0.001, n > 14.

### Intracellular nucleotide availabilities limit cellular fitness

To determine whether the decreased growth of GC-rich cells also translates into a decreased competitive fitness relative to AT-rich cells, coculture experiments were performed with randomly chosen pairs of AT-and GC-rich strains. As expected, GC-rich strains were readily outcompeted by AT-rich strains (Fig. 4): In all 32 replicate populations, a strong decrease in the frequency of GC-rich cells relative to AT-rich cells was observed within the first two days of the experiment. Already after two days, GC-rich strains went extinct in 50% of all experimental populations. At the end of the experiment (i.e. after eight days), GC-rich strains were present in only two out of 32 populations (i.e. 6 %), suggesting a strongly reduced fitness of GC-rich cells relative to AT-rich cells.

**Fig.4.**
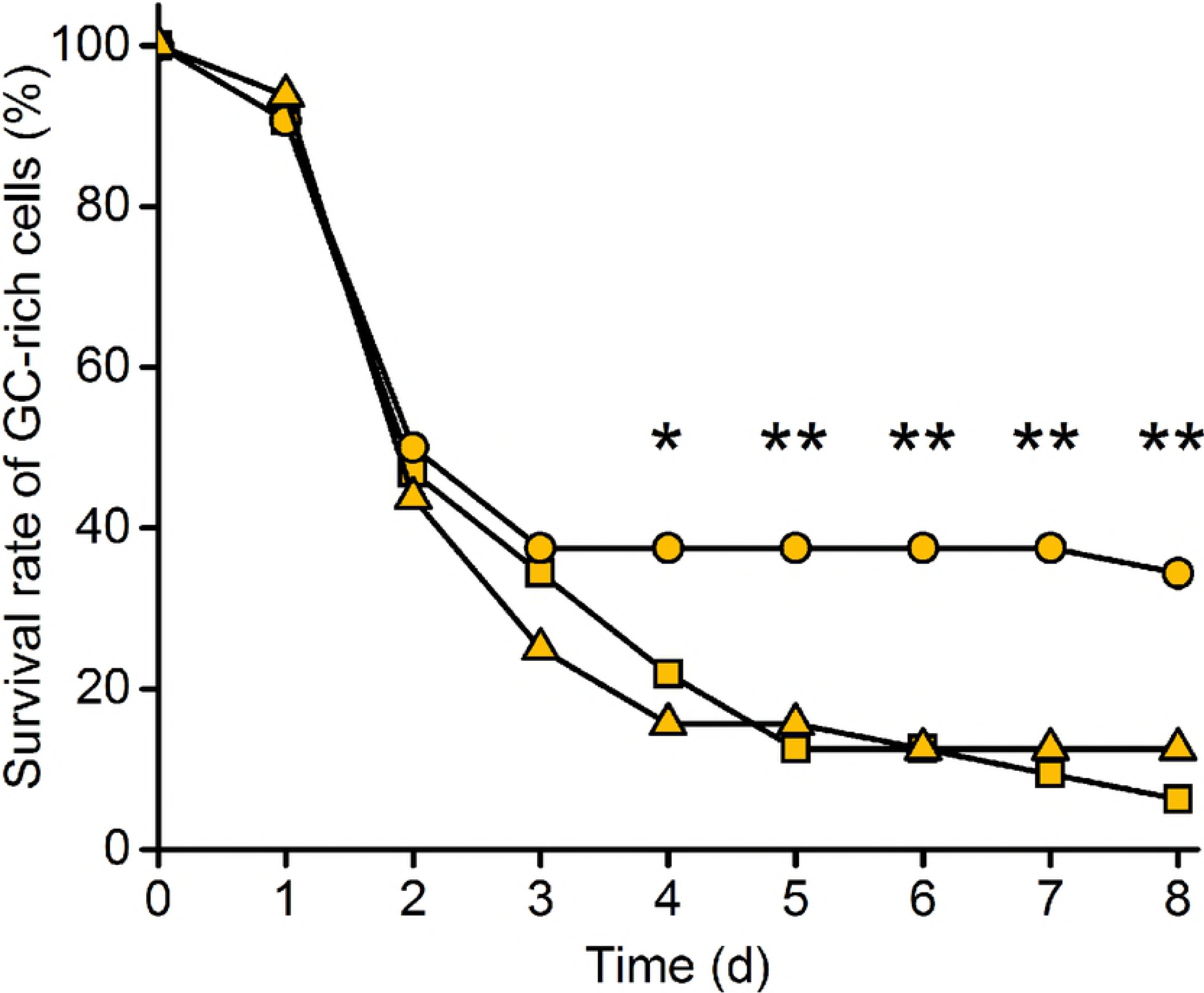
GC-supplementation partially rescues fitness of cells with GC-rich plasmids. Eight pairwise cocultures of cells harboring AT-and GC-rich pJet plasmids were co-inoculated (1:1 ratio) and serially propagated on a daily basis. Strains were supplied with AT or GC deoxyribonucleosides (100 μM per nucleoside). Survival rates of GC-rich *E. coli* strains were calculated by scoring the number of populations, in which GC-rich strains were still present relative to the total number of populations within each treatment (n = 32). Each of the eight pairwise combination of strains was replicated four times. Treatment groups comprise the unsupplemented control (triangles), supply with AT-nucleosides (squares), and supply with GC-nucleosides (circles). Asterisks indicate survival rates that were significantly different from the unsupplemented control group. False discovery rate-corrected Wilcoxon signed ranks-test: ** P < 0.01, * P < 0.05, n = 32.

Two main explanations can account for this pattern. First, the growth of GC-rich cells could have been limited by the availability of G+C nucleotides within cells, while the cells containing AT-rich plasmids were less strongly affected. This hypothesis would be in line with the idea that the molecular composition of extrachromosomal genetic elements is strongly affected by natural selection. Second, other properties of GC-rich DNA in general could limit the growth of cells containing GC-rich plasmids. For example, an increased stability of GC-rich DNA could raise the chances for stable secondary structures that might hamper DNA replication (19) and thus slow down growth. These two effects can be distinguished in a competition experiment between AT-rich and GC-rich cells, in which cocultures are supplemented with either A+T or G+C nucleotides. If the observed decrease in fitness of GC-rich cells was truly due to a lack of G+C nucleotides within cells, then providing GC-rich cells with G+C nucleotides should enhance their growth more than supplementation with A+T. In contrast, if another mechanism such as e.g. the formation of secondary structures applies, nucleotide supplementation should not differentially affect the growth of AT-and GC-rich cells.

For this experiment, nucleosides instead of nucleotides were used, since *E. coli* can take up nucleosides, but not nucleotides (20). When A+T nucleosides were externally provided, the survival rate of GC-rich cells was not significantly different from the one of the unsupplemented control group (Fig. 4), indicating that these nucleosides do not generally limit cellular growth. When G+C nucleosides were added to the growing cultures, the decline of GC-rich strains observed within the first three days of the experiment was as fast as in the untreated control group. However, from day three onwards, GC-rich cells continued to survive in ~35% of the cocultures, which represents a significant increase over both the A+T-supplemented and the untreated control group (Fig. 4). This finding indicates indeed that the low availability of G+C nucleotides limited the growth of GC-rich cells, thereby corroborating the hypothesis that the availability of nucleotides in the host cytoplasm plays a key role in shaping the GC-content of extrachromosomal genetic elements.

### The cost of extrachromosomal elements depends on the GC-content of the host chromosome

The finding that AT-rich plasmids impose a lower metabolic burden on host cells than GC-rich plasmids was obtained using *E. coli* as host, whose chromosome has a GC-content of ~ 50%. However, the intracellular availability of nucleotides likely depends on the base composition of the cell’s chromosome, because the biosynthetic machinery of a cell is expected to have evolved in a way that it produces the required building block metabolites in optimal amounts. As a consequence, G+C nucleotides should be more abundant than A+T nucleotides in species with high genomic GC-contents, thus rendering GC-rich plasmids less costly than AT-rich plasmids.

In order to test this hypothesis, eight bacterial species with a genomic GC-content ranging from 40% GC *(Acinetobacter baylyi* ADP1) to 68% GC *(Azospirillum brasilense*Tarrand) were transformed with one randomly chosen pair of AT-and GC-rich pBAV plasmids. In this context, it should be noted that the GC-rich plasmid was characterized by a net GC-content of 51% (AT-plasmid: 33% GC) and was thus enriched in GC-nucleotides when compared to the chromosomes of species with low to intermediate GC-contents. However, it displayed a slightly lower GC-content when compared to the genome of GC-rich species such as *Azospirillum brasilense* and *Xanthomonas campestris.* Subsequently, the fitness of different host cells carrying the GC-rich versus AT-rich plasmids was quantified spectrophometrically as before. Indeed, the results of these experiments revealed that the burden imposed by these two plasmids depended on the genomic GC-content of the bacterial host: when the host chromosome was more AT-rich, the fitness of cells containing the AT-rich plasmid was higher than the one of cells containing the GC-rich plasmid (Fig. 5). In contrast, when host cells with a more GC-rich chromosome were considered, the fitness of cells containing the GC-rich plasmid was increased relative to cells containing the AT-rich plasmid (Fig. 5). Thus, the molecular composition of the host’s genome strongly affected the fitness cost imposed by extrachromosomal genetic elements.

**Fig.5.**
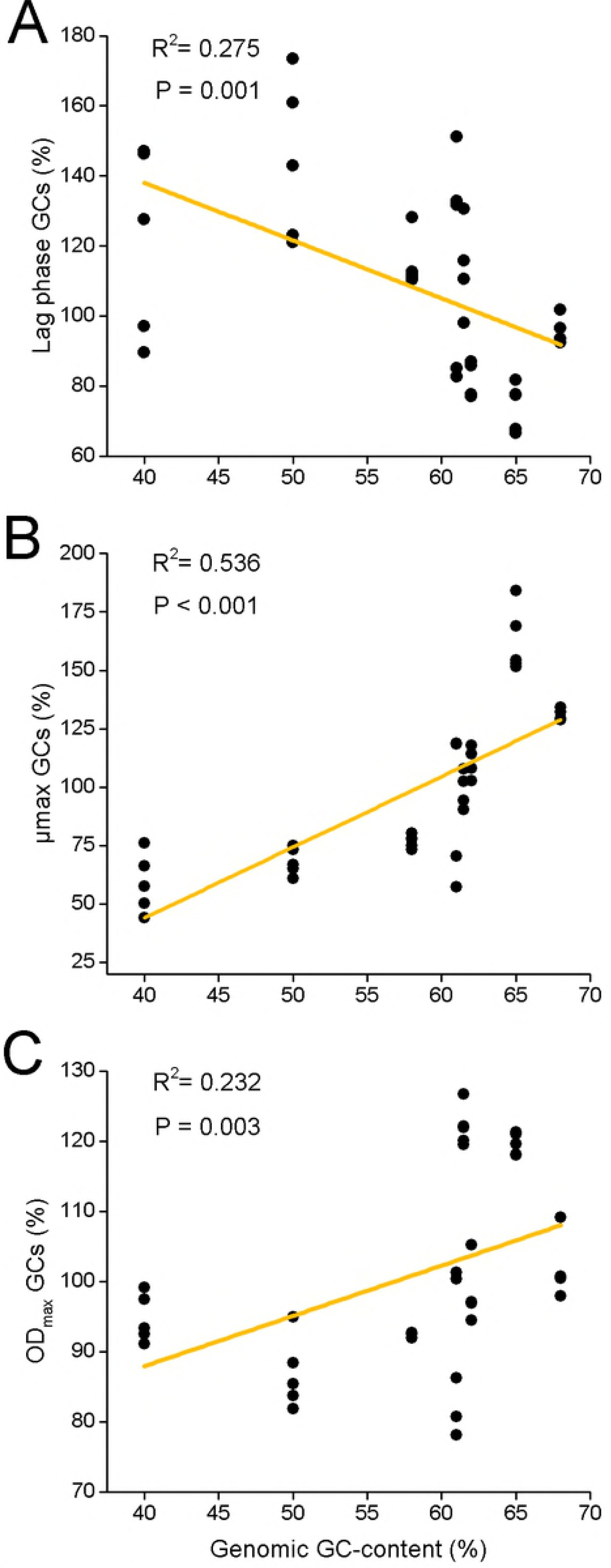
The fitness cost of plasmids depends on the GC-content of both plasmid and host genome. Growth experiments of eight bacterial species that differ in their genomic GC-content and which harboured one AT-or GC-rich pBAV plasmid were performed in minimal medium. Fitness parameters of cells with GC-rich plasmids were calculated relative to the respective strain harbouring the AT-rich plasmid. (**A**) Lag-phase, (**B**) maximum growth rate, and (**C**) maximum optical density reached after 24 h. Linear Regression analysis, n = 5 in all cases.

In addition, plasmid copy number measurements revealed that AT-rich plasmids were generally more abundant in species with low and intermediate genomic GC-contents (Fig. S1). However, in the two species with the highest GC-contents (i.e. *Azospirillum brasilense* and *Xanthomonas campestris),* GC-rich plasmids were present in higher copy numbers than the AT-rich plasmids.

Taken together, GC-rich plasmids imposed higher fitness costs than AT-rich plasmids on species with low to intermediate genomic GC-contents, while AT-rich plasmids were more costly for bacterial species with higher genomic GC-contents.

## Discussion

The DNA of intracellular, host-dependent elements such as bacterial endosymbionts and plasmids is generally more AT-rich than the DNA of their host’s genome. In the case of bacterial endosymbionts, this pattern is commonly thought to be due to neutral evolutionary processes, such as genetic drift or a mutational bias. However, here we provide strong experimental evidence that selective advantages can contribute to this pattern. By experimentally manipulating the GC-content of plasmids and quantifying the resulting fitness consequences for the corresponding bacterial host, we show that the fitness cost of plasmids strongly depended on the nucleotide composition of both the plasmid and the host’s genome. Specifically, when the genome of the host cell was characterized by intermediate to high A+T content, GC-rich plasmids were more costly (Figs. 2 and 5) and present in a lower copy number than AT-rich plasmids (Figs. 3 and S1). In contrast, when the host chromosome was enriched in G+C, GC-rich plasmids were less costly (Fig. 5) and present in a higher copy number than AT-rich plasmids (Fig. S1). Supplementation experiments confirmed that the observed fitness effects were indeed due to limiting pools of the corresponding nucleotides and not resulting from the GC-content of the introduced sequences *per-se* (Fig. 4).

The continuous synthesis of nucleotides is crucial for DNA replication in all dividing cells. Under optimal conditions, *E. coli* cells can divide every 20 minutes, while the replication of the chromosome takes about 40 minutes (21). To overcome this problem, multiple chromosomal copies are simultaneously generated, such that replication can keep up with the speed of cell division (22). Interestingly, not the activity of the polymerase, but the nucleotide biosynthesis required for DNA replication seems to be limiting growth, as evidenced by the observation that the shortage of a single nucleotide drastically decreases growth (23). Moreover, up-regulation of the ribonucleotide reductase (RNR) in yeast, which is responsible for the synthesis of dNTPs, increases the speed of the replication fork (24). Both studies show that nucleotide synthesis is the rate-limiting step for DNA synthesis and hence also growth. By linking the availability of nucleotides to cellular fitness, these studies support the main findings reported here.

Our results are consistent with an evolutionary scenario, in which the intracellular availability of nucleotides (A+T versus G+C) depends on the genomic nucleotide composition of the bacterial cell. Over evolutionary time, a bacterial cell likely establishes an equilibrium, in which the biochemical machinery that produces all four nucleotides is tailored to meet the cell’s requirements. This includes not only the nucleotides that are required for DNA replication, but also those that are needed to produce RNA, signalling molecules (e.g. ppGpp, cAMP), or coenzymes (e.g. ATP). In addition, energetic and stoichiometric parameters are likely to affect nucleotide production rates. For example, the biosynthetic cost to produce A+T nucleotides is less than the energy that is required to biosynthesize the same amounts of G+C (4). An extrachromosomal genetic element that now enters such a cellular system disturbs this equilibrium by withdrawing nucleotides from intracellular pools to enable its own replication. By doing so, plasmids (and likely also intracellular bacterial endosymbionts) incur a cost to the hosting cell that depends on both the nucleotide availability in the host’s cytoplasm and the amount and identity of nucleotides it consumes.

In cells of *Escherichia coli,* whose genome is characterized by a mean AT-content of ~ 50%, ATP is the most abundant nucleotide (3.5 mM ATP, 2.0 mM UTP, 1.9 mM GTP, and 1.2 mM CTP under exponential growth, see (10)). This is likely because of the dual function of ATP, which is not only used for RNA and DNA synthesis, but also plays a key role for transferring energy within cells. Unfortunately, to the best of our knowledge, no other study exists to date that quantified cytoplasmic nucleotide concentrations in other bacterial species, especially those that feature higher genomic GC-contents. Nevertheless, it appears reasonable to assume that cells with a higher genomic GC-content should also have an increased demand for G+C nucleotides including both ribo-and deoxyribonucleotides for RNA-and DNA-biosynthesis, respectively. This would imply higher cytoplasmic G+C levels and thus render the consumption of G+C nucleotides by GC-rich plasmids potentially less detrimental than in host cells with high genomic AT content and thus, low cytoplasmic G+C. The observation that plasmid copy numbers of AT-rich plasmids were higher in species with low to intermediate genomic GC-contents, but decreased in species with more GC-rich genomes (Fig. S1), is in line with this hypothesis.

Previous work that compared the GC-content of plasmids with the genomes of their corresponding bacterial and archaeal hosts revealed that on average, the GC-content of plasmids was lower than the GC-content of their host’s genome (i.e. between 3% and 10%; see (4) and (5)). In light of the results presented in our study, this finding suggests that A+T nucleotides are likely always more abundant in the cytoplasm of bacterial hosts and/ or cheaper to produce than G+C nucleotides. Two fundamentally different processes could cause the observed increased AT-content of plasmids relative to the genome of their bacterial host. First, plasmids could evolve towards increased AT-contents relative to their host’s chromosome. Second, plasmids with an increased relative AT-content might be more successful in establishing in new host cells via routes of horizontal gene transfer (i.e. via conjugation or transformation). However, how would selection operate to favour extrachromosomal genetic elements with an increased A+T content? In principle, selection can operate on two different levels. First, several intracellular elements that differ in their genomic composition can compete against each other. If selective advantages are sufficiently strong, elements with an increased AT-content should outcompete elements with a lower AT-content, thus resulting in a globally increased AT-content of the entire population of intracellular elements (symbiont-level selection). Alternatively, selection can act on the level of the host (host-level selection). In this case, host individuals that contain more AT-rich elements are evolutionarily fitter than hosts containing more GC-rich intracellular elements. As a consequence of the resulting competition, hosts that contain more AT-rich elements will survive and reproduce with a higher chance, thus favouring AT-rich elements on the level of the host population in the long-run. For the given experimental set-up, our results demonstrate that host-level selection can be strong and result in an almost complete elimination of GC-rich plasmids within a few days (Fig. 4). Unfortunately, it was not possible to test whether AT-rich plasmids could outcompete GC-rich plasmids within a given host cell, as plasmids using the same mode of replication cannot coexist within the same cell (i.e. when they belong to the same plasmid incompatibility group, see (25))

Our results not only help to understand the GC-content variation in extrachromosomal genetic elements such as plasmids and viruses, but have also significant ramifications for endosymbiotic bacteria. Similar to the interaction between bacteria and their plasmids, host-dependent bacterial cells regularly feature genomes with drastically increased AT-contents relative to the DNA of their host cell. In addition, many bacterial endosymbionts have lost the genes for an autonomous biosynthesis of all four nucleotides (26, 27). Hence, to maintain a sufficient nucleotide-supply, cells require uptake mechanisms that allow them to import nucleotides from the host’s cytoplasm. Indeed, uptake systems for nucleotide triphosphates in intracellular bacteria have been previously identified for *Rickettsia* and *Chlamydia* (28, 29), which are also known to lack specific genes essential for nucleotide biosynthesis pathways.

Bacterial endosymbionts (30), and in fact the majority of prokaryotic and eukaryotic organisms (31, 32), display a characteristic mutational bias that generally increases the genomic A + T content. In addition, newly established endosymbionts sometimes show an unexpectedly large number of polymerase slippage events that preferentially eliminate G+C-rich repetitive sequences, thus also biasing the endosymbiont’s genome towards an increased A + T content (33). Finally, population bottlenecks that occur frequently when populations of bacterial endosymbionts are vertically transmitted from parent to offspring host, result in random assortment of bacterial genotypes that can lead to the fixation of AT-rich symbiont populations within hosts. In the early onset of an endosymbiotic interaction, all of the abovementioned processes are likely selectively neutral. However, at some point, host individuals that harbour symbionts with increased AT-contents will display a higher fitness than hosts that contain more GC-rich symbionts, particularly given the large number of endosymbiont cells in an individual host that amplify the costs associated with nucleotide requirements of the symbiont population. Due to this fitness difference, host-level selection should favour hosts with metabolically ‘cheap’ AT-rich symbionts. We thus believe that the evolution of AT-rich endosymbionts is likely a combination of both neutral processes such as mutational bias/ genetic drift and natural selection.

Taken together, our results provide strong experimental support for the hypothesis that the availability of nucleotides represents a significant evolutionary force that shapes the base composition of host-dependent, extra-chromosomal elements such as plasmids and likely also endosymbiotic bacteria. This interpretation is at odds with the widely-held view of drift as being the sole explanation for the AT-bias observed in the genomes of host-restricted bacteria. While our study adds an important new facet to this on-going discussion, it is most likely a combination of multiple factors that determines the nucleobase composition of bacterial genomes.

## Materials and Methods

### Identification of AT-and GC-rich sequences

Eight AT-and GC-rich stretches of 1 kb in size each were identified from the AT-rich genome of *Arabidopsis thaliana* Col-0 (genome version TAIR9 v171 obtained from the Plant Genomic Database) and GC-rich genome of *Chlamydomonas reinhardtii* wild type 137 C (assembly and annotation v4 obtained from DOE Joint Genome Institute). Both annotated genomes were imported into Geneious (version 6.1.8, Biomatters, New Zealand) (34) that was used to identify AT-and GC-rich DNA stretches, respectively. Importantly, sequences were selected such that no promoters, start codons, or regulatory elements were present. Moreover, sequences containing simple sequence repeats in a total length of more than 30 bp were excluded to avoid the possible formation of stem-loop structures. PCR primers (Table S1) were designed using the software Primer 3 (35) and ordered from Metabion International AG (Martinsried, Germany).

### Amplification of AT-and GC-rich sequences

Genomic DNA of *A. thaliana* Col-0 was extracted following the method of Allen *et al.*(36) and of *C. reinhardtii* wild type 137 C using the protocol described by (37). AT-rich DNA was amplified by PCR using Phusion HiFi, Polymerase (Fermentas/ Thermo Fisher Scientific, Waltham, Massachusetts, US) following the manufacturer’s protocol. PCR program: 98 °C 1 min, 30x: 98 °C 15 s, T_m_ primer 15 s, 68 °C 40 s. Elongation temperatures were decreased to 68 °C according to Su *et al.* (38), since no PCR product was observed at 72 °C. GC-rich DNA was amplified from *C. reinhardtii* using Kapa2G Robust Polymerase (Peqlab; Erlangen, Germany) following the manufacturer’s recommendations for GC-rich DNA. PCR program: 95 °C 5 min, 30x: 95 °C 15 s, T _m_primer 5 s, 72 °C 40 s. PCR products were purified by gel electrophoresis (1% agarose) using the NucleoSpin Extract II gel and PCR clean-up Kit (Macherey-Nagel GmbH & Co. KG, Duren, Germany).

### Construction of AT-and GC-rich plasmids

Two plasmid backbones were used for the insertion of AT-and GC-rich DNA. The first backbone was pJet1.2/blunt (Thermo Fisher Scientific), a commercially available, high copy number plasmid of small size. The plasmid carries a pMBI* origin of replication and encodes a beta-lactamase *(bla)* that confers resistance to ampicillin, which was used to select for plasmid-containing cells. AT-and GC-rich sequences were inserted into the blunt-end multiple cloning site of pJet using the pJet1.2/blunt Cloning Kit (Thermo Fisher Scientific) lacking the P_lacUV5_ promoter. Plasmids were transformed into chemically competent *E. coli* TOP10 cells (Invitrogen, Thermo Fisher Scientific) using the heat shock method (39). Transformed colonies were screened for the respective insert using the Colony Fast-Screen Kit (Epicentre; Madison, Wisconsin, USA) following the manufacturer’s instructions. Plasmids of selected transformants were sequenced at MWG Eurofins (Ebersberg, Germany).

To validate the experimental results and exclude plasmid-specific effects, all AT-and GC-rich sequences were additionally inserted into a second, high copy number plasmid, a modified pBAV1kT5-gfp (17) (ordered from Addgene https://www.addgene.org/; Cambridge, Massachusetts, US). This plasmid uses a different replication system (i.e. repA-mediated replication) and encodes a different selectable marker, aminoglycoside-3’-phosphotransferase *(aph(3’)),* which confers resistance to the antibiotic kanamycin. The gene encoding the green fluorescent protein present on the plasmid was not needed for this study and hence removed by digesting the plasmid with *NotI* (Thermo Fisher Scientific). Blunt ends were generated and clean-up was carried out as described above. AT-and GC-rich sequences were inserted into the same position. The resulting plasmids were transformed into chemically competent *E. coli* TOP10 cells. Transformants were sequenced in order to validate loss of the T5-gfp cassette and successful insertion of the AT-and GC-rich sequences. In the main text, the modified plasmid lacking *T5-gfp* is denoted as pBAV instead of pBAV1Kt5-gfp.

### Bacterial strains

All AT-rich plasmids were transformed into *E. coli* BW25113 Ara-(40), whereas GC-rich plasmids were transformed into *E. coli* BW25113 Ara+ (41), respectively that were made chemically competent using the rubidium chloride method (39). The *Δara* mutation renders the strain unable to catabolize arabinose. Both strains can be phenotypically distinguished when plated on tetrazolium arabinose indicator plates, on which *E. coli*BW25113 (Ara+) forms white and BW25113 *Δara* (Ara-) red colonies (42, 43). The arabinose marker is selectively neutral under the cultivation conditions used in this study (independent-samples t-test: P > 0.05, n = 8). This phenotypic marker was used to distinguish both strains when grown in coculture.

### Culture conditions and growth kinetics

All experiments were performed in M9 minimal medium (44), which was complemented with 2 mM MgSO_4_, 0.1 mM CaCl_2_, and 5 g l^-1^ Glucose (Sigma, St. Louis, Missouri, USA). For pBAV-containing strains, 0.25% Casamino acids (Sigma) were added to promote growth. Precultures were prepared by streaking genotypes from glycerol stocks on Lysogeny Broth (LB, Sigma) agar plates (Thermo Fisher Scientific), which were incubated overnight (16 h) at 37 °C. Subsequently, single colonies were picked and inoculated into 0.8 ml of M9 medium in a 96-deepwell plate (Eppendorf, Hamburg, Germany), which was then incubated overnight at 37 °C under shaking conditions. To ensure plasmid maintenance, 100 μg ml^-1^ ampicillin or 50 μg ml^-1^ kanamycin (Sigma) were always added to the culture media for pJet-and pBAV-harbouring strains, respectively. Next, optical densities (OD determined at a wavelength of 600 nm) of all precultures were measured in a microwell platereader (Spectramax 250, Molecular Devices; Sunnyvale, USA) using a 96-well plate (Nunc, Fisher Scientific GmbH; Schwerte, Germany) with a culture volume of 200 μl. The OD_600nm_ of each culture was adjusted to 0.001. Growth kinetic assays were performed in the same instrument. Using a culture volume of 200 μl, growth was measured as absorbance at 600 nm every 5 min at 37 °C for 24 h. Cultures were shaken for 3 min after each and 15 s prior to each measurement. Fitness-relevant growth parameters (i.e. lag phase, maximum growth rate, and maximal OD_600nm_) were calculated using Magellan 7.1 SP 1 software (Magellan Software GmbH; Dortmund, Germany).

### Plasmid copy number determination

Plasmid copy numbers were determined using quantitative real-time PCR (qPCR) following a previously described method (45). For this, monocultures of cells were harvested after 24 h of growth. Plasmid copy numbers were determined by calculating both the total number of chromosomal and plasmid copies in each sample. Chromosome copy numbers were determined using a primer pair targeting the single copy gene *dxs* (1-deoxy-D-xylulose-5-phosphate synthase). For total plasmid numbers per sample, primer pairs targeting the respective antibiotic resistance gene were used (i.e. either *bla* on pJet or *aph(3’)* on pBAV; genes and primer details see Table S2). Bacterial cultures from both monocultures were diluted ~1:100 and used for qPCR. QPCR was performed using the Brilliant III Ultra-Fast SYBR Green QPCR Master Mix (Agilent Technologies; Santa Clara, US) in a BioRad CFX96 thermocycler (Hercules, California, USA) according to the manufacturer’s instructions. PCR program: 10 min 95 °C, 40x: 30 s 95 °C, 20 s 61 °C, 30 s 72 °C. Standard curves were prepared by 10-fold dilutions of both isolated plasmids and bacterial cells (R^2^ of all standard curves: > 0.99). Plasmid numbers per ng plasmid DNA template were calculated using an online calculator (http://cels.uri.edu/gsc/cndna.html, Andrew Staroscik, Genomics & Sequencing Center, University of Rhode Island, Kingston, Rhode Island, USA). Cell numbers of each standard curve sample were measured using a CyFlow Space flow cytometer (Partec, Görlitz, Germany), for which cells were stained with SYBR Green (Sigma) following the manufacturer’s protocol. Average plasmid copy numbers per cell were calculated from the respective standard curves (R^2^ of all standard curves > 0.99) by dividing total plasmid numbers by the total number of cells.

### Competitive fitness and nucleoside feeding experiments

For coculture experiments, one AT-rich strain was paired with a GC-rich strain, respectively (all harbouring pJet plasmids). Eight combinations were chosen randomly (i.e. Fig. 1: AT1-GC1, AT2-GC2, etc.) and each combination replicated 4 times (n = 32). The OD_600nm_ of all precultures was adjusted to 0.0005, resulting in a final OD_600nm_ of 0.001 after mixing of cocultures. Fitness experiments were performed in a 96-deepwell plate (Nunc) with a culture volume of 0.8 ml. Cocultures were incubated at 37 °C under shaking conditions (220 rpm). 0.8 μl (1:1,000 dilution) of all cocultures were transferred daily into fresh M9 minimal medium. Every day, serial dilutions of all cultures were plated on TA agar plates to distinguish Ara+ (AT-rich) and Ara-strains (GC-rich) of *E. coli*BW25113 using the arabinose utilization marker as described above.

Moreover, cocultures were supplied with nucleosides to test if the decrease in growth of GC-rich cells can be explained by their increased demand for GC nucleotides. No nucleotide transport systems are known for *E. coli,* however, two nucleoside transport systems have been described: The nucleoside permeases NupG and NupC use the proton motive force to import nucleosides (20). Hence, deoxyribonucleosides were used for this experiment. Either 2’-deoxyadenosine and thymidine or 2’-deoxyguanosine and 2’-deoxycytidine (Sigma) were added to the growth medium at a final concentration of 100 μM per nucleoside.

### Transformation of bacterial species with different GC-contents

One AT-and one GC-rich pBAV plasmid, i.e. pBAV-AT01 and pBAV-GC02, were randomly chosen to be introduced into other bacterial species differing in their genomic GC-contents. PJet could not be used for this purpose as it does not replicate in species other than *E. coli.* In contrast, pBAV has been shown to be a broad host range plasmid replicating in many other bacterial species (17). The following species were used: *Acinetobacter baylyi* ADP1 (40% GC), *Serratia entomophila* (DSM 12358) (58% GC), *Pseudomonas protegens* (61% GC), *Pseudomonas putida* (62% GC), *Arthrobacter aurescens* (DSM 20116) (61.5% GC), *Xanthomonas campestris* (DSM 3586) (65% GC), and *Azospirillum brasilense* (DSM 1690) (68% GC). All strains were tested to be Kanamycin (50 μg/ml) sensitive. Both plasmids were introduced in electrocompetent cells of the above listed species using a MicroPulser Electroporator (Bio-Rad, Hercules, California, US) with the following settings: 25 μF, 200 mA, and 2.5 kV using 70 μl of electrocompetent cells and 100-150 ng plasmid DNA. Colonies obtained were grown in LB medium supplemented with 50 μg ml^-1^ kanamycin. Plasmid isolation was performed as described previously and plasmids were sequenced using plasmid-specific primers targeting the AT/GC-rich insert.

### Growth experiments and plasmid copy number determination using bacterial species with different GC-contents

All strains were grown in M9 minimal medium containing glucose, sucrose, and malate (glucose and sucrose: 5 g l^-1^ each, malate: 2 g l^-1^) as carbon source, as well as 2 mM MgSO_4_, 0.1 mM CaCl_2_, 45 μM FeSO_4_, 0.5 mg ml^-1^ NaMO_4_, and 0.01 mg ml^-1^ Biotin (Sigma) at 28°C and 220 rpm for 24 h. Growth experiments were performed as mentioned above using a Spectramax plate reader. Plasmid copy numbers of different bacterial species were determined by measuring plasmid numbers via quantitative RealTime PCR as described earlier. However, all cell numbers were quantified by Flow Cytometry instead of qPCR, since using *dxs* specific primers did not result in DNA amplification in most of the species, either due to an altered sequence or absence of the gene. Thus, plasmid copy numbers were calculated as plasmid number per cell count (see above).

### Data analysis

Experiments were performed using plasmids that contained one of eight different AT-or GC-rich inserts. To reduce the impact of sequence-specific effects, data were analysed by treating AT-rich and GC-rich strains as replicates. In monoculture experiments, fitness-relevant parameters of AT-and GC-rich cells were statistically compared by two-sample independent t-tests. In coculture experiments, survival rates of strains harbouring GC-rich plasmids were calculated by scoring the number of populations, in which the strains were present relative to the total number of populations within each treatment (n = 32). Strains were considered to be extinct, if the number of colony forming units (CFUs) decreased below 2% of the total CFU counts. Survival rates between treatments were compared by performing Wilcoxon signed ranks tests. False discovery rate (FDR) was applied to P-values to correct for multiple testing (46). Linear regression analyses were performed to correlate growth parameters of species harbouring AT/GC-rich plasmids with the species’ GC-content. All statistical analyses were performed using SPSS Software (version 17.0, SPSS Inc., Chicago, IL, USA) and R Studio (Boston, USA) (47).

### Author contributions

CK, MK, and AKD conceived the study. All authors designed the experiments, AKD conducted all experiments, analysed the data, and wrote the first draft of the manuscript. All authors amended the manuscript. Authors declare no competing interests.

## Acknowledgements

We thank the Experimental Ecology and Evolution Group, the Insect Symbiosis Group, the Department of Bioorganic Chemistry as well as Günter Theißen for helpful discussion and Colin Dale and Rahul Raghavan for critically reading an earlier version of the manuscript. Moreover, help by Eva Limpinsel with generating some of the strains and support by Wilhelm Boland is gratefully acknowledged.

## Supporting Information Legends Figures

### Figures

**Fig. S1. Plasmid copy number of GC-rich plasmids compared to AT-rich plasmids is increased in bacterial species with GC-rich genomes.** Copy numbers of GC-rich relative to AT-rich plasmids in the same bacterial species are displayed. Plasmid copy number per cell equivalent was assessed by quantitative real-time PCR and flow cytometry of all bacterial species after 24 h of growth. Asterisks denote significant deviations from equal copy numbers of AT-and GC-rich plasmids in the same host environment. One sample t-test: *** P < 0.001,** P < 0.01, * P < 0.05, n = 5.

### Tables

**Table S1. Non-coding AT-and GC-rich sequences used in this study.** AT-rich sequences (AT01-08) were amplified from the genome of *Arabidopsis thaliana*(chromosome *4),* whereas GC-rich sequences (GC01-GC08) were amplified from *Chlamydomonas reinhardtii* (+chromosome 1, ‘chromosome 2).

**Table S2. Primer pairs used for quantitative real-time PCR to determine the copy number of pJet and pBAV plasmids relative to the copy number of the chromosome.**

